# Key role of amino acid repeat expansions in the functional diversification of duplicated transcription factors

**DOI:** 10.1101/014910

**Authors:** Núria Radó-Trilla, Krisztina Arató, Cinta Pegueroles, Alicia Raya, Susana de la Luna, M.Mar Albà

**Affiliations:** Evolutionary Genomics Group, Research Programme on Biomedical Informatics (GRIB), Hospital del Mar Research Institute (IMIM), Dr. Aiguader 88 08003 Barcelona, Spain.; Universitat Pompeu Fabra (UPF), Dr. Aiguader 88 08003 Barcelona, Spain.; Centre for Genomic Regulation (CRG), Dr. Aiguader 88, 08003 Barcelona, Spain.; Centro de Investigación Biomèdica en Red en Enfermedades Raras (CIBERER), Spain.; Institució Catalana de Recerca i Estudis Avançats (ICREA), Pg. Lluís Companys 23, 08010 Barcelona, Spain.

**Author notes:** Corresponding authors contact information: **M. Mar Albà** ICREA Research Professor, Research Programme on Biomedical Informatics (GRIB), Hospital del Mar Research Institute (IMIM) - Dr. Aiguader 88, 08003 Barcelona, Spain.; **Susana de la Luna** ICREA Research Professor, Gene Regulation, Stem Cells and Cancer Programme, Centre for Genomic Regulation (CRG) - Dr. Aiguader 88, 08003 Barcelona, Spain.

**Keywords:** gene duplication, low complexity region (LCR), transcription factor, PHOX2B, polyalanine

## Abstract

The high regulatory complexity of vertebrates has been related to two closely spaced whole genome duplications (2R-WGD) that occurred before the divergence of the major vertebrate groups. Following these events, many developmental transcription factors (TFs) were retained in multiple copies and subsequently specialized in diverse functions, whereas others reverted to their singleton state. TFs are known to be generally rich in amino acid repeats or low-complexity regions (LCRs), such as polyalanine or polyglutamine runs, which can evolve rapidly and potentially influence the transcriptional activity of the protein. Here we test the hypothesis that LCRs have played a major role in the diversification of TF gene duplicates. We find that nearly half of the TF gene families (107 out of 237) originated during the 2R-WGD contain LCRs, compared to only a small percentage of the non-duplicated TF genes used as a control (15 out of 115). At the individual gene level, we observe that twice as many duplicated TFs have gained LCRs as non-duplicated TFs. In addition, duplicated TFs preferentially accumulate certain LCR types, the most prominent of which are alanine repeats. We experimentally test the role of alanine-rich LCRs in two different TF gene families, PHOX2A/PHOX2B and LHX2/LHX9. In both cases, the presence of the alanine-rich LCR in one of the copies (PHOX2B and LHX2) significantly increases the capacity of the TF to activate transcription. Taken together, the results provide strong evidence that LCRs are important driving forces of evolutionary change in duplicated genes.

## Introduction

Gene duplication is a major mechanism for the emergence of novel gene functions (Ohno 1970). Duplicated genes may arise from local genomic duplication events affecting just one or a few genes, or from whole genome duplications (WGDs). It has been observed that after a WGD there is preferential retention of duplicated genes that are in dosage balance, including many transcription factors and developmental regulatory proteins (Birchler et al. 2005; Freeling and Thomas 2006; Edger and Pires 2009). At later evolutionary stages, these factors diverge and acquire new functions (Escriva et al. 2006; Lynch et al. 2008). Many human transcription factors (TFs) encoded in the human genome date from the two rounds of whole genome duplication (2R-WGD) that occurred in early vertebrate evolution (Ohno et al. 1968; Lundin 1993; Dehal and Boore 2005; Blomme et al. 2006). The 2R-WGD increased the opportunities for new regulatory pathways to arise, contributing to the formation of complex structures such as the brain and the circulatory system (Wagner 2008; Huminiecki and Heldin 2010).

Many studies have reported an increase in the number of amino acid substitutions after gene duplication, which is consistent with relaxed purifying selection and/or gain of new functions in one or both gene copies (see for example Zhang et al. 2003; Scannell and Wolfe 2008; Han et al. 2009; Pegueroles et al. 2013; Pich I Roselló and Kondrashov 2014). Other works have examined the divergence of expression patterns of gene duplicates, which appears to be relatively independent of amino acid sequence change (Wagner 2000; Makova and Li 2003; Farré and Albà 2010).

However, there is a lack of studies on amino acid repeat expansions after gene duplication. These events often correspond to insertions or deletions in the alignments and they become invisible when evolutionary rates are computed. Over time, perfect amino acid tandem repeats formed by triplet slippage tend to accumulate amino acid substitutions and become cryptic repeats or simple sequences (Green and Wang 1994; Huntley and Golding 2000; Albà et al. 2002; Simon and Hancock 2009). Both perfect and cryptic repeats, collectively called low-complexity regions (LCRs), can be detected on the basis of sequence compositional biases by specific programs, such as SEG (Wootton and Federhen 1996). It has been noted that LCRs are especially abundant in TFs compared to other functional classes (Karlin et al. 2002; Albà and Guigó 2004; Faux et al. 2005; Huntley and Clark 2007).

Several lines of evidence point to LCRs as a source of functional innovation. First, tracts of repeated proline, glutamine, or alanine fused to a DNA-binding domain can change the capacity of a protein to activate transcription (Gerber et al. 1994; Sauer et al. 1995; Janody et al. 2001; Galant and Carroll 2002; Brown et al. 2005). Second, histidine repeats can mediate the targeting of the protein to nuclear speckles, which are subnuclear structures for the storage of splicing factors and TFs (Salichs et al. 2009). Third, the uncontrolled expansion of glutamine repeats has been associated with a number of neurodegenerative diseases in humans (Gatchel and Zoghbi 2005), whereas mutations resulting in abnormally long alanine repeats cause several developmental disorders (Messaed and Rouleau 2009). Fourth, changes in the length of amino acid repeats have been associated with morphological changes in dog breeds (Fondon and Garner 2004). Finally, although repeats can expand rapidly by triplet slippage, there is also evidence that they are often preserved by purifying selection once they reach a certain length (Mularoni et al. 2010).

Here, we investigate the role of LCRs in the functional diversification of human duplicated TFs formed during the 2R-WGD. Using this set of families ensures that the number of genes is sufficiently large and homogeneous for statistical analysis. In addition, we can experimentally test the functional consequences of LCRs in TFs using well-established transcriptional activity assays. We find that there is a very significant enrichment of LCR in duplicated TFs when compared to non-duplicated TFs. The results of the experiments performed in two different gene families, PHOX2A/PHOX2B and LHX2/LHX9, illustrate how the gain of a novel LCR in one of the gene copies alters the capacity of the protein to activate transcription, connecting the presence of the LCR to a gain of functional activity.

## Results

### Increased gain of low complexity regions (LCRs) in duplicated transcription factors (TFs)

In order to investigate the role of LCRs in the functional diversification of gene duplicates, we obtained a set of 550 human TFs that had been retained in duplicated form after the vertebrate 2R-WGD. For this, we used paralogy information from Ensembl (Vilella et al. 2009; Flicek et al. 2013) as well as phylogenetic tree reconstructions based on homologous sequences from diverse vertebrate species (see Material and Methods). The TFs belonged to 237 different gene families with 2, 3, or, in two cases, 4 gene members in the human genome (Figure 1, duplicated transcription factors; supplementary tables S1 and S2, Supplementary Material online). The scarcity of gene families of size 4 concurs with the observation that many duplicated genomic segments were lost following the 2R-WGD (Friedman and Hughes 2003; Dehal and Boore 2005). For comparison, we also obtained a set of 115 human non-duplicated TF genes (Figure 1, single-copy transcription factors; supplementary table S3, Supplementary Material online). These genes presumably reverted to their single copy gene state soon after the 2R-WGD.

**Figure 1.**
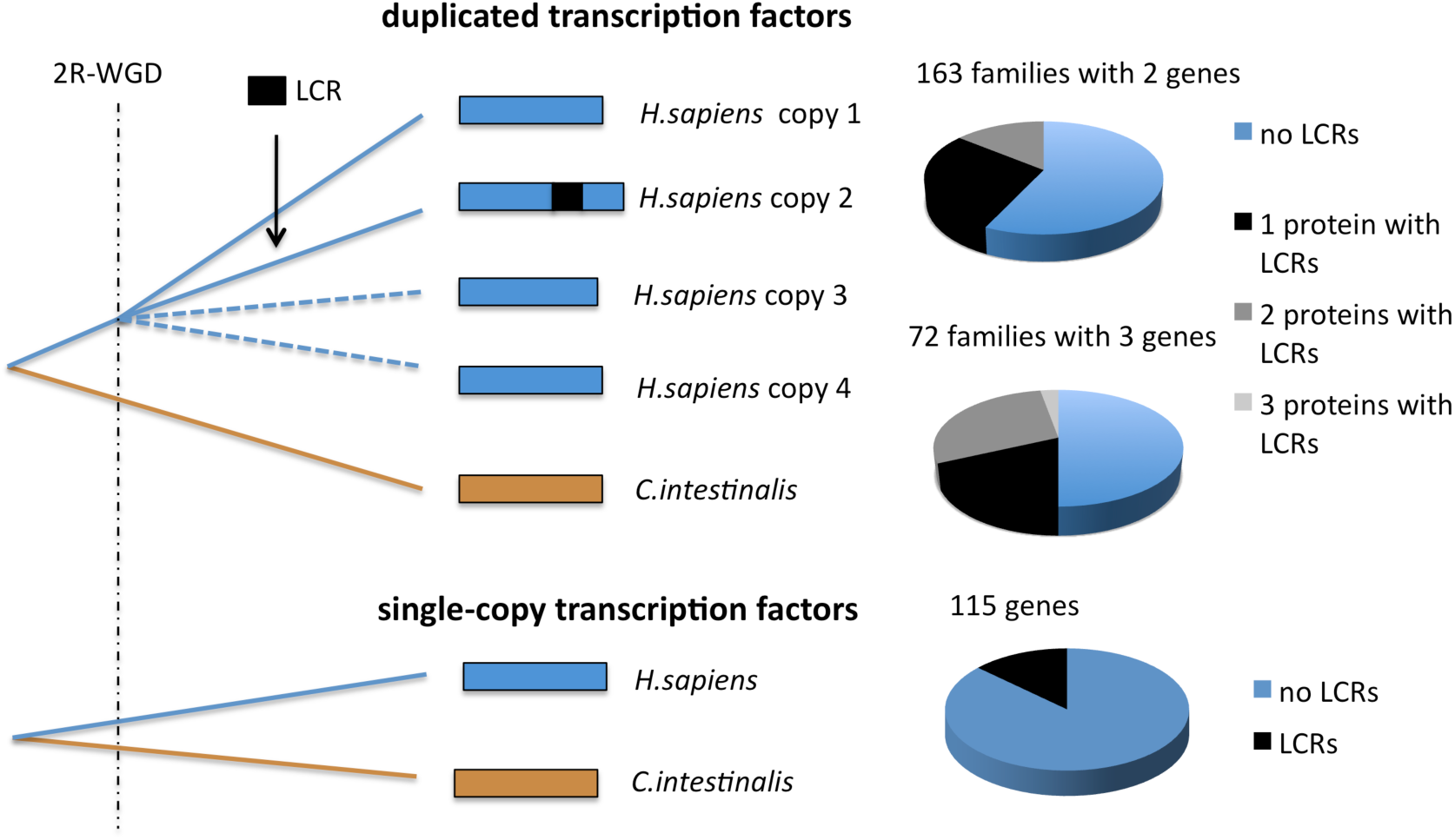
Diagram showing the gene family datasets studied and the number of proteins with LCRs in each of them. The number of duplicated gene families was 237: 163 had 2 members, 72 had 3 members, and only 2 had 4 members (one protein in this latter subset had LCRs, not shown in the Figure). The total number of proteins with repeats in the duplicated gene set was 155. The number of single copy genes in the non-duplicated datset was 115, of which 15 had LCRs.

We identified LCRs in TFs with the SEG algorithm (Wootton and Federhen 1996), using parameter settings that favor the detection of highly repetitive sequences (see Material and Methods). LCRs ranged between 9 and 59 amino acids in size, with an average of 22 residues. 107 of the 237 gene families (45%) contained LCRs in one or more gene copies (Figure 1). This set comprised 222 LCRs in 155 different duplicated proteins out of the 550 analyzed (supplementary tables S4 and S5, Supplementary Material online). In contrast, we only found 20 LCRs in 15 single copy proteins out of the 115 analyzed (Figure 1, supplementary table S6, Supplementary Material online). Therefore, the fraction of proteins with LCRs in the duplicated gene set was about twice as large as the fraction in the non-duplicated gene set (28% versus 13%, respectively, Chi-square test p=0.009; supplementary table S7, Supplementary Material online).

After careful examination of all multiple sequence alignments of the gene families with LCRs, we could only identify 8 cases (out of 222) in which the LCR conservation pattern was consistent with the existence of a repeat before the 2RWGD. Similarly, using *Ciona intestinalis* orthologues we could only detect three putative cases of chordate ancestral repeats in our data set. Therefore, we concluded that the vast majority of the LCRs identified (>95%) had been formed after the 2R-WGD.

The most common scenario was gain of LCRs in one member of the gene family, but we also found several cases in which LCRs had been gained in an independent manner in several family members. One illustrative example is the gene family comprising *POU4F1, POU4F2* and, *POU4F3.* Each of these genes contains two exons each and encodes a highly conserved POU domain at the protein C-terminus. We identified LCRs in the POU4F1 and POU4F2 proteins, but not in POU4F3. POU4F1 contained one LCR enriched in alanines and another one in glycines. POU4F2 had a histidine-rich tract, previously shown to be required for nuclear speckle targeting (Salichs et al. 2009), and another long LCR containing several serine and glycine runs. The LCRs from the two proteins mapped to different exons and were clearly unrelated to each other.

We identified 62 LCRs that were highly conserved in different vertebrate lineages and that had probably originated in the period between the 2R-WGD and the separation of the bony fishes, approximately 450 to 420 million years ago (Dehal and Boore 2005). In other cases the incomplete conservation of the LCR suggested a more recent time of origination, such as the alanine-rich LCRs in PHOX2B and LHX2, which apparently formed after the amphibian split and which are discussed in more detail below.

### Overrepresentation of alanine-rich LCRs in duplicated TFs

Next, we investigated the amino acid composition of the LCRs in the duplicated and non-duplicated gene sets. Most LCRs were highly enriched in one or at most two amino acids (supplementary tables S4 to S6, Supplementary Material online). For this reason we decided to label each LCR according to the most frequent amino acid or, in those cases in which two different amino acids were similarly abundant, the two most frequent amino acids (17% of LCRs). The most commonly found amino acids in the non-duplicated gene set were glutamic acid and proline (6 and 4 cases, respectively). In contrast, in the duplicated gene set the most abundant amino acid was alanine, accounting for 50 of the 222 LCRs (22.5%, in 42 proteins)(Figure 2). Remarkably, we only detected one alanine-rich LCR in the non-duplicated gene set (1 out of 20 LCRs; 5%).

**Figure 2.**
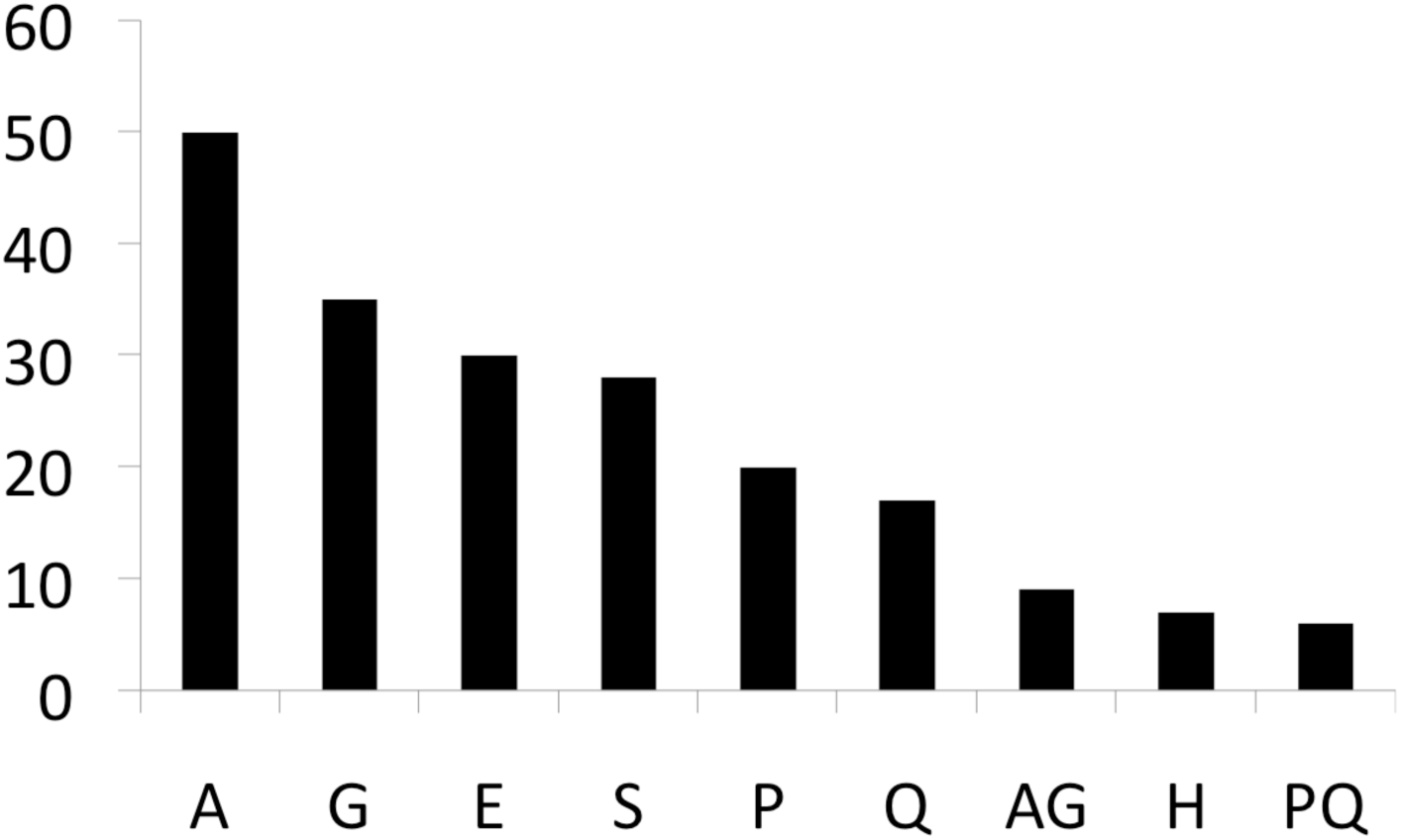
Number of LCRs annotated by the most abundant amino acid(s) in the duplicated gene set. The list has been shortened to LCR types occurring more than 5 times.

The list of duplicated proteins with alanine-rich LCRs included several disease-associated genes (supplementary table S8, Supplementary Material online).Noteworthy, we identified two genes in which expansions of the alanine tract beyond the normal range result in an incorrect functioning of the protein in humans: Paired-like homeobox 2B (*PHOX2B*) and Aristaless related homeobox (*ARX*). The LCR in the PHOX2B protein contains a perfect run of 20 alanines that, when expanded to 25-29 residues, causes central hypoventilation syndrome, an alteration in the development of the autonomic nervous system (Amiel et al. 2003; Trochet et al. 2005). The ARX sequence includes four alanine repeats; short expansions beyond the wild type range in the first and second repeats are found in individuals with X-linked mental retardation and in several other X-linked developmental disorders (Shoubridge et al. 2010). Given these examples, it seems possible that mutations increasing the size of the alanine repeats in other genes of the set may be associated with developmental disorders of yet unknown origin.

### The effect of alanine-rich LCRs on transcriptional activation

To evaluate the functional impact of the LCRs gained after gene duplication, we examined in detail two transcription factor families: PHOX2A/PHOX2B and LHX2/LHX9 (LIM homeobox 2/9). In both families one of the gene copies had gained an alanine-rich LCR (PHOX2B and LHX2). In PHOX2B the LCR included a perfect alanine tract of size 20, whereas in LHX2 there was a perfect repeat composed of 10 alanines. Whereas the polyalanine tract in PHO2B has been associated with central hypoventilation syndrome, no previous data exists on the possible functionality of the alanine repeat in LHX2.

We experimentally tested the transcriptional activity of different gene constructs, including the two paralogous proteins and a deletion mutant lacking the LCR. We employed one-hybrid assays on a reporter gene in which the DNA binding sites were placed on a minimal promoter structure (containing only a TATA-box) to reduce promoter-specific effects. Moreover, we used a heterologous DNA binding domain (DBD) to tether the effector protein to the reporter DNA to avoid any confounding effect due to possible differential DNA binding affinities of the different family members.

## PHOX2A/PHOX2B

The paralogous genes *PHOX2A* and *PHOX2B* encode transcription factors that play key roles in different developmental processes of vertebrates. Both genes are expressed in peripheral and central noradrenergic neurons and neural crest derivatives, with differences in the onset and extent of expression during development (Pattyn et al. 1997; Amiel et al. 2003). The association of each paralogue to a specific functional role is further complicated by the fact that *PHOX2B* activates *PHOX2A* as observed in gain-of-function experiments (Flora et al. 2001). However, both paralogues are not functionally redundant because *PHOX2A* cannot always replace *PHOX2B in vivo* (Coppola et al. 2005). *PHOX2A* is critical for the development of noradrenergic neurons and specific motor neuron nuclei (Pattyn et al. 1997). *PHOX2B* is essential for the development of the central and peripheral autonomic nervous systems and it is a master regulator for the differentiation of the visceral nervous system (Pattyn et al. 1999; Dauger et al. 2003).

The alignment of the two human paralogous proteins shows that the homeodomain, which functions as the DNA-binding domain (Benfante et al. 2007), is extremely well conserved between the two copies and that most differences accumulate in a region at the C-terminal end that has several amino acid repetitions; the most conspicuous of which is an alanine-rich LCR only present in *PHOX2B* (Figure 3).

**Figure 3.**
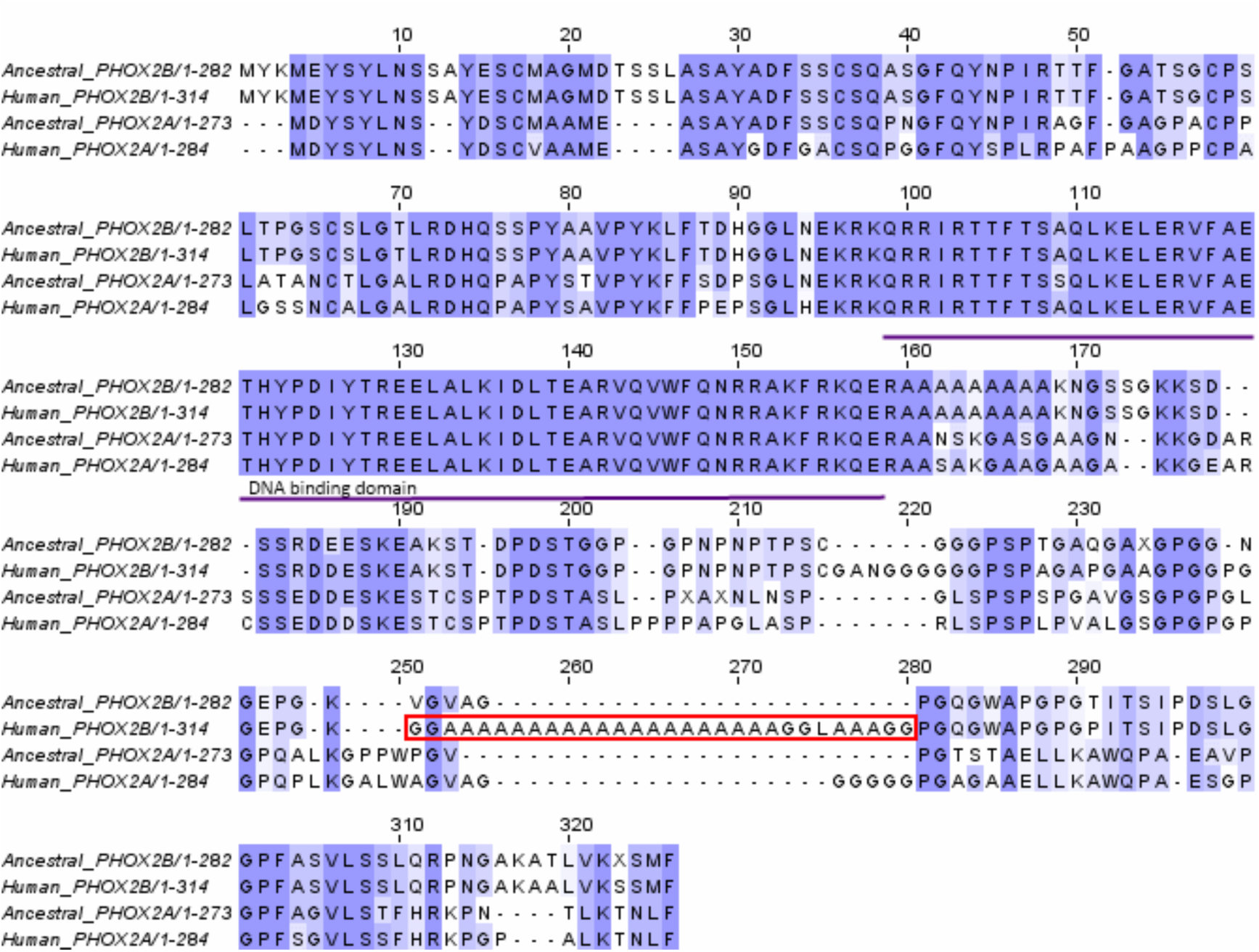
Alignment of human PHOX2B and PHOX2A proteins, together with the corresponding ancestral, non-LCR containing, proteins. The low-complexity region (LCR) identified in the human PHOX2B by SEG is framed. Sequence reconstruction was performed by maximum likelihood (see Material and Methods for more details). We employed sequences from Xenopus, chicken, cow, rat, mouse, macaque, and human. Zebrafish sequences were not included because the examination of sequences from the different species clearly indicated that the LCR had been gained after the separation from the amphibians.

Analysis of orthologous sequences from several vertebrates indicated that the alanine-rich LCR in PHOX2B had probably originated in an ancestral *Amniota.* We combined sequence information from *Xenopus*, chicken, human, macaque, mouse, rat and cow (supplementary table S9, Supplementary Material online) to deduce the most likely ancestral sequence at a time when the LCR was not yet present using a maximum likelihood approach (Yang 2007). We followed the same procedure for the PHOX2A protein. The sequence alignment of ancestral and modern proteins shows that the reconstructed ancestral PHOX2B is about 10% shorter than the human PHOX2B and lacks the expanded alanine repeat (Figure 3).

We generated expression plasmids to produce the complete PHOX2A and PHOX2B open reading frames which were N-terminally fused to Gal4-DBD (Figure 4A). We also constructed a plasmid to express a mutated version of PHOX2B that lacked the alanine-rich LCR (PHOX2BΔLCR). The three fusion proteins accumulated to a similar extent when over-expressed in HeLa cells, as shown by the results of Western blot analysis with an antibody against the Gal4-DBD (Figure 4B). Immunofluorescence analysis revealed that the proteins were located in the nuclei of the transfected cells, suggesting that the repeat was not affecting subcellular localization (Figure 4C). In reporter assays, PHOX2B showed a dose-dependent transcriptional activity over the basal Ga4-DBD levels, which was not detected in transfections with PHOX2A (Figure 4D; supplementary table S10, Supplementary Material online for the results of three experimental replicates). In PHOX2BΔLCR the capacity to activate transcription was strongly impaired with relation to the wild type PHOX2B, providing strong evidence that the alanine-rich LCR is an activator domain. These results support the idea that the formation of the LCR had a strong impact in the functional divergence of the two gene copies.

**Figure 4.**
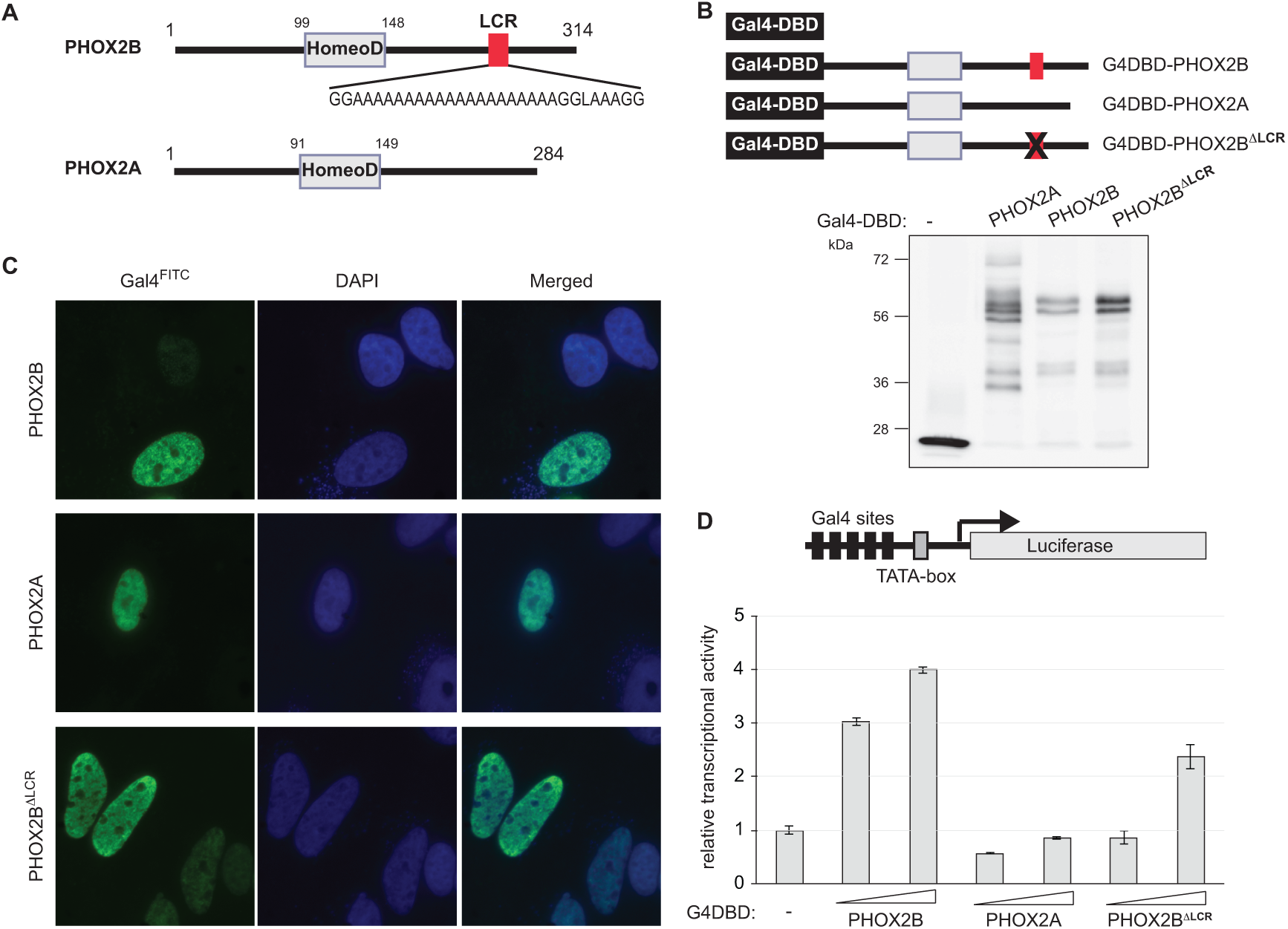
**A)** Schematic representation of PHOX2A and PHOX2B. HomeoD: homeodomain; LCR: low-complexity region. **B)** Total cell extracts from cells transfected with equal amounts of the plasmids to express the indicated Gal4DBD-fusion proteins (as indicated in the scheme) were analyzed by Western blot with anti-Gal4-DBD to show the expression levels of the fusion proteins. **C)** The subcellular localization of the indicated PHOX2-fusion proteins (as in B) was analyzed by indirect immunofluorescence with anti-Gal4-DBD (left panels) in HeLa transfected cells. Nuclei were counterstained with DAPI (middle panels). **D)** One-hybrid reporter assays using a 5xGal4 sites/luciferase reporter and increasing amounts (150 ng, 450 ng) of expression plasmids to the effector proteins indicated. The graph represents the transcriptional activation of Gal4-DBD fusions to PHOX2A, PHOX2B wild-type or to the PHOX2B LCR deletion mutant, as indicated, relative to that of unfused Gal4-DBD arbitrarily set as 1. Error bars represent standard deviations of triplicated transfected plates. The experiment shown is representative of 3 performed.

## LHX2/LHX9

The *LHX2* and *LHX9* genes belong to the LIM homeobox (LHX) family, a group of proteins which perform important roles in tissue-specific differentiation and body patterning during development in both vertebrates and invertebrates (Kadrmas and Beckerle 2004). In particular, both *LHX2* and *LHX9* are involved in the development of brain, limb, and eyes, as well as in hematopoiesis (Srivastava et al. 2010). Consistent with the formation of *LHX2* and *LHX9* by gene duplication during the 2R-WGD, the two paralogoues are present in different vertebrate lineages but there is only one orthologous gene, named *Apterous*, in the *Drosophila* genus. *Apterous* shares key functional properties with *LHX2/LHX9* genes during development (Hobert and Westphal 2000).

The LHX2 and LHX9 proteins contain two LIM domains, a type of zinc-finger known to mediate protein-protein interactions (Matthews et al. 2009), and a helix-turn helix forming a homeodomain that binds to specific DNA sequences in the promoters of target genes (Figure 5). The paralogous protein LHX2 additionally contains an alanine-rich LCR that is highly conserved in mammals. We reconstructed ancestral sequences for the two paralogues using a similar approach than with the PHOX2A/PHOX2B pair (supplementary table S11, Supplementary Material online). The alignment indicated that the modern and ancestral LHX2 proteins are vey similar except for the alanine-rich LCR (Figure 5).

**Figure 5.**
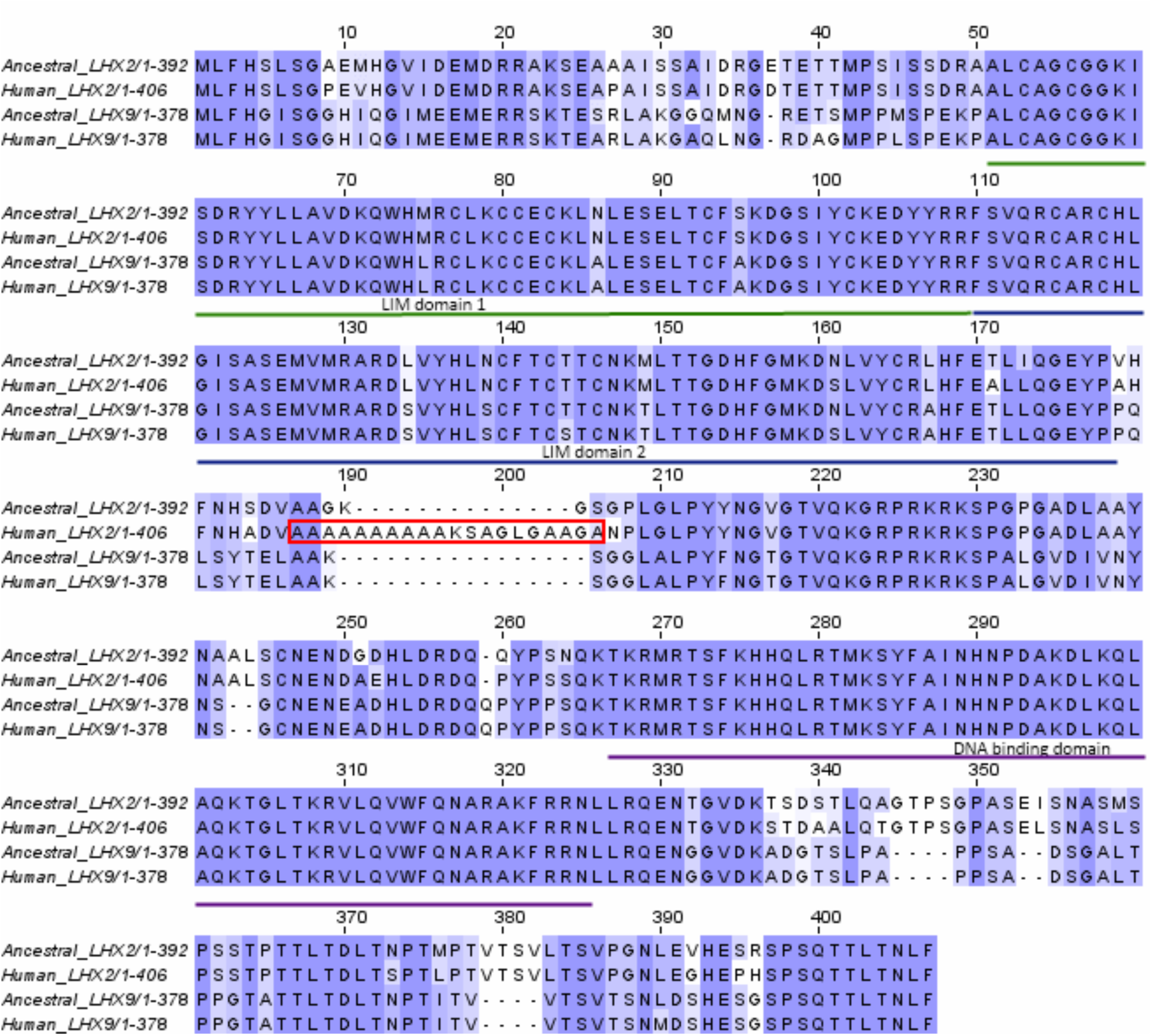
Alignment of human LHX2 and LHX9 proteins, together with the corresponding ancestral non-LCR containing proteins. The low-complexity region (LCR) identified in the human LHX2 protein by SEG is framed. Sequence reconstruction was performed by maximum likelihood (see Methods for more details). We employed sequences from Xenopus, chicken, cow, rat, mouse, macaque, and human. Zebrafish sequences were not included because the examination of sequences from the various species clearly indicated that the LCR had been gained after the separation from the amphibians.

We experimentally tested the transcriptional activity of the two gene copies and the specific effect of the LHX2 alanine-rich LCR using the one hybrid assay (Figure 6). The two paralogues and the LCR deletion mutant of LHX2 were expressed at similar levels (Figure 7B) and the subcellular distribution was indistinguishable (Figure 6C). LHX2 behaved as an activator in this experimental setting, but LHX9 did not show any significant difference with respect to the control containing only the DNA-binding domain (Figure 6D; supplementary table S12, Supplementary Material online for the results of three experimental replicates). The LHX2 activity was severely impaired when the LCR was deleted. The results clearly link the formation of the LCR with the gain of a novel functionality in LHX2.

**Figure 6.**
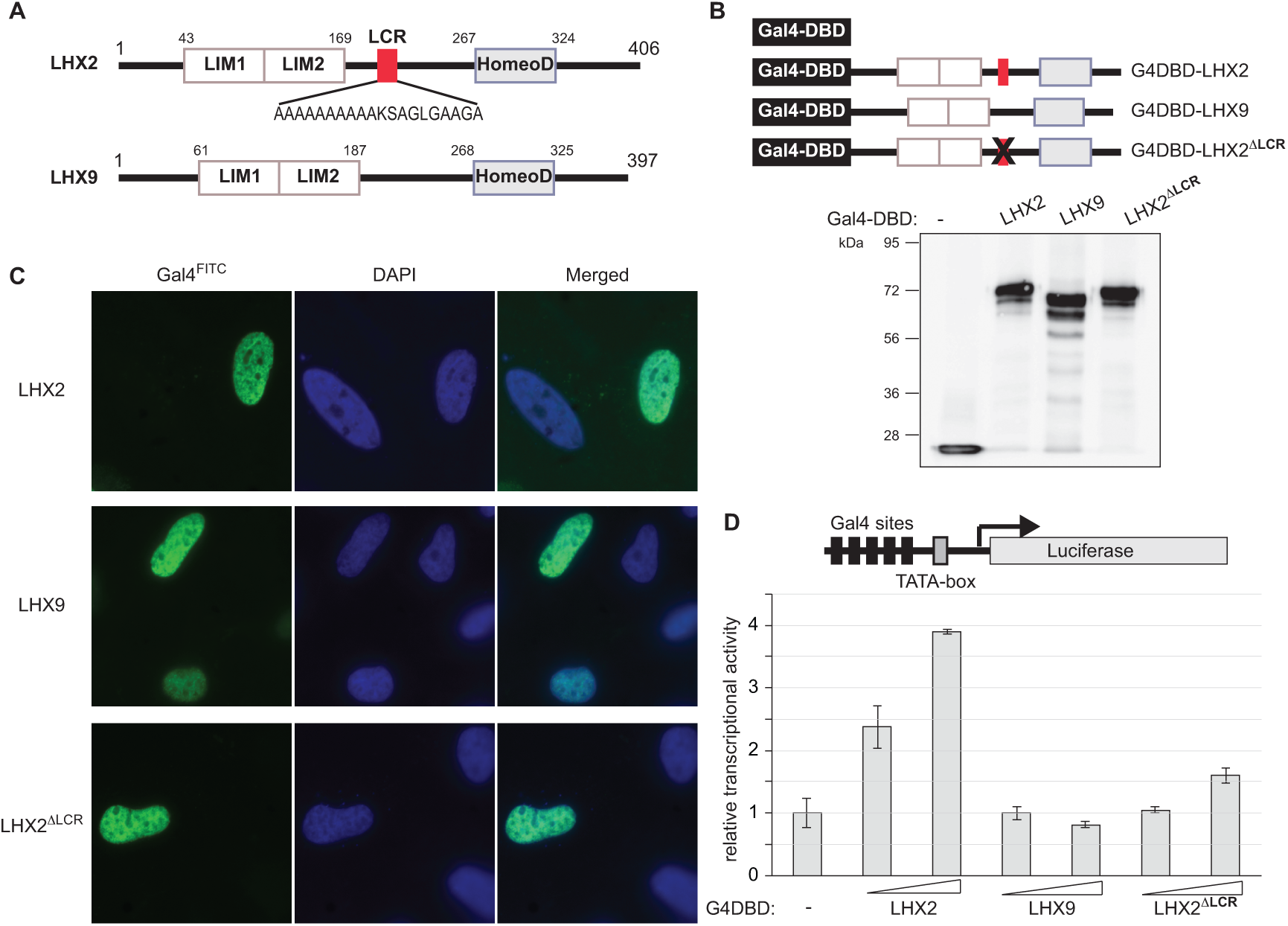
**A)** Schematic representation of LHX2 and LHX9. HomeoD: homeodomain; LCR: low-complexity region; LIM: LIM domain. **B)** Total cell extracts from cells expressing transfected with equal amounts of the plasmids to express the indicated Gal4DBD-fusion proteins (as indicated in the scheme) were analyzed by Western blot with anti-Gal4-DBD to show the expression levels of the fusion proteins. **C)** The subcellular localization of the indicated fusion proteins (as in B) was analyzed by indirect immunofluorescence with anti-Gal4-DBD in transfected HeLa cells (left panels). Nuclei were counterstained with DAPI (middle panels). **D)** One-hybrid reporter assays using a 5xGal4 sites/luciferase reporter and increasing amounts (50 ng, 150 ng) of expression plasmids to the effector proteins indicated. The graph represents the transcriptional activation of Gal4-DBD fusions to LHX9, LHX2 wild-type or to the LHX9-LCR deletion mutant, as indicated, relative to that of unfused Gal4-DBD arbitrarily set as 1. Error bars represent standard deviations of triplicated transfected plates. The experiment shown is representative of 3 performed.

## Discussion

The evolution of vertebrates is intimately related to the two rounds of whole genome duplication (2R-WGD) that occurred before the diversification of the group (Lundin 1993; Dehal and Boore 2005; Cañestro et al. 2013). Many developmental genes were retained after the 2R-WGD and subsequently gained new functions, a process known as neofunctionalization. One classic example is the retinoic acid receptor (RAR) family, which includes RAR alpha, RAR beta, and RAR gamma. Whereas RAR beta maintained its original function, the other two paralogues accumulated amino acid substitutions that modified their capacity to bind different retinoid compounds (Escriva et al. 2006).

Here, we turned our attention to modifications due to gain of LCRs. Except for a few examples, including a number of proteins containing histidine repeats (Salichs et al. 2009), the incidence of LCRs in the evolution of gene duplicates has not been examined to our knowledge. We found that nearly half the TF gene families (107 out of 237) originated at the 2R-WGD contained LCRs, and that the number of duplicated genes that had gained LCRs was significantly larger than the equivalent number of non-duplicated genes. In some cases we even found that several members of the same family had gained different LCRs.

In general, the LCRs corresponded to long gaps when aligned to the other gene copies, consistent with the action of replication slippage in expanding the repeat. In the absence of functionality, repeats will tend to accumulate amino acid substitutions and degenerate. However, the sudden disappearance of an LCR,albeit possible, appears to be a rare event (Radó-Trilla and Albà 2012). Similarly, LCRs are more frequently found associated with insertions than with deletions in multiple sequence alignments of mammalian orthologous proteins (Laurie et al. 2012). Taken together, these observations point to an important role of repeat expansions in increasing protein length and complexity.

Our bioinformatics pipeline identified 50 different alanine-rich LCRs, ranging in size from 9 to 31 amino acids, in the duplicated TFs. Glycine-rich LCRs followed in order of abundance (30 gains) and there were additional 9 LCRs that were enriched in both alanine and glycine. Repeats of these two amino acids appear to expand with relatively ease in vertebrate proteins in general (Nakachi et al. 1997; Radó-Trilla and Albà 2012). The reasons are yet unclear but several factors, such as high slippage propensity of GC-rich triplets (Ellegren 2000), low amino acid synthesis cost, reduced toxicity, or high adaptive value of the runs formed by these amino acids, may all contribute to this bias.

Experiments performed in the gene families *PHOX2A/PHOX2B* and *LHX2/LHX9* linked the presence of alanine-rich LCRs to neofunctionalization. In both cases we found that the LCR significantly increased the ability of the protein to activate transcription. Although the precise molecular mechanisms of this activation remain to be determined, one possibility is that it is driven by the interaction with coactivator subunits of the basal transcription complex TFIID, as shown for the alanine-rich region of the *Drosophila* TF Bicoid (Sauer et al. 1995).

The impact of alanine repeats in the regulatory activity of PHOX2B, and more specifically the consequences of abnormal expansions of these tracts for human health, have been the focus of several studies. In humans, the heterozygotic expansion of the 20-alanine stretch in PHOX2B to 25-29 residues leads to congenital central hypoventilation syndrome (CCHS; OMIM 603851), a rare disorder defined by a failure of the autonomic control of breathing (Amiel et al. 2003). It has been reported that several PHOX2B mutants with expansion of the alanine repeat suffer an alteration in the conformation of the PHOX2B protein that disrupts normal protein function and impairs the ability of the PHOX2B to regulate the transcription of its target genes (Bachetti et al. 2005; Trochet et al. 2005; Di Lascio et al. 2013). In addition, PHOX2B nonsense mutations leading to truncated proteins that lack the C-terminus, which contains the alanine repeat, have been found in neuroblastoma patients (Raabe et al. 2008). PHOX2B polyalanine expanded mutants show a significant reduction in their capacity to activate transcription when assayed *in vitro* (Adachi et al. 2000), suggesting that when the tract becomes abnormally long it has a negative impact on transcriptional activity. Interestingly, expansion of the alanine tract from 15 to 25 residues in the ZIC2 protein, which would mimic the mutation found in holoprosencephaly patients, also results in near complete loss of transcriptional activation (Brown et al. 2005).

In addition to the link between transcriptional activation and the presence of a poly-alanine tract, the evolutionary conservation of the LCR might be associated with other protein activities. For example, deletion of the alanine repeat in FOXL2 results in intranuclear protein aggregation (Moumné et al. 2005). In addition, the combination of alanine repeats with other LCR types may lead to complex functional outcomes. It has been reported that alanine repeats can exert an inhibitory effect over glutamine repeats, which are strong activators on their own (Janody et al. 2001; Galant and Carroll 2002). Analysis of GO terms in our set of duplicated TFs with yielded a similar number of genes annotated as “positive regulation of transcription” than of genes annotated as “negative regulation of transcription” among those containing alanine-rich LCRs (supplementary table S5, Supplementary Material online), suggesting that these repeats may influence transcription in very different contexts.

In summary, the results of the present study indicate that the rate of gain of LCRs in duplicated genes increases with respect to single copy genes. As shown for alanine repeats, this can result in protein neofunctionalization. This process is likely to have been shaped by selection, contributing to increase the complexity of regulatory networks in vertebrates.

## Material and Methods

### Sequence datasets and identification of gene families

We selected human genes whose molecular function in the Gene Ontology database matched “transcription factor” in Ensembl v.64/66, using BioMart (Flicek et al. 2013). We annotated TFs with an activator or repressor role using the Gene Ontology (GO) terms “positive regulation of transcription” and “negative regulation of transcription” (Ashburner et al. 2000). We used the paralogous gene dating information in Ensembl Compara (Vilella et al. 2009) to extract human transcription factor families whose duplication time was consistent with the two rounds of whole genome duplication (2R-WGD) at the base of the vertebrates before the formation of jawed vertebrates (Euteleostomi paralogy type).

For each gene family member, orthologous sequences from *Mus musculus* (mouse), *Gallus gallus* (chicken), and *Danio rerio* (zebrafish) were retrieved from Ensembl. We obtained 539 proteins in mouse, 441 proteins in chicken, and 635 proteins in zebrafish, denoting some secondary loses and duplications (in fishes due to an additional WGD) with respect to the human dataset. We also obtained orthologous proteins in *C. intestinalis* whenever available (141 *C. intestinalis* proteins). We used these sequences to generate multiple protein alignments with T-Coffee (Di Tommaso et al. 2011). We subsequently built maximum likelihood based trees with PhyML v.2.4.4 (Guindon et al. 2005) and only accepted tree topologies congruent with the hypothesis of 2R-WGD, with no more than two duplication events in the ancestral vertebrate branch and no subsequent duplications in the lineage leading to humans. The final set of duplicated proteins consisted of 237 transcription factor protein families: 163 families with two human gene copies, 72 families with three human gene copies, and 2 families with four human gene copies. The total number of human genes within these families was 550 (average 2.32 genes/family).

A set of 115 non-duplicated human transcription factors was also obtained from Ensembl for comparative purposes. These proteins corresponded to transcription factors for which no paralogues had been retained after the 2R-WGD.

### Identification and characterization of low complexity regions (LCRs)

We identified sequences corresponding to low complexity regions (LCRs) using the program SEG (Wootton and Federhen 1996). This program divides sequences into contrasting segments of low sequence complexity, amino acid repeats, and segments of high sequence complexity. The low sequence complexity segments are identified because they depart strongly from a random residue composition. We used the parameters window=15, K_1_=1.5, and K_2_=1.8, which identify highly repetitive regions.

The amino acid repeats were annotated according to the most frequently occurring amino acid in the sequence identified by SEG. When the frequency of the second amino acid was more than half the frequency of the most abundant one (17% of the repeats), the two most frequently occurring amino acids were considered in the annotation of the repeats. In general these cases corresponded to the presence of several single amino acid repeats of different composition next to each other.

We used an in-house Perl script to identify the position of all LCRs on the previously generated multiple sequence alignments. Conserved LCRs were defined as those that overlapped in the alignment and were enriched in the same amino acid.

### Reconstruction of ancestral sequences

Vertebrate ancestral protein sequences, including ancestral LHX2, LHX9, PHOX2A, and PHOX2B, were reconstructed using Codeml (parameter RateAncestor = 1) in PAML v4.4 (Yang 2007). All multiple sequence alignments included orthologous proteins from *Homo sapiens, Macaca mulata, M. musculus, Rattus norvegicus, Bos taurus, Xenopus tropicalis, D. rerio*, and in the case of LHX2 and LHX9, also *G. gallus* (supplementary tables S9 and S11, Supplementary Material online). Ancestral sequence reconstruction was based on alignments obtained with Prank +F to increase the accuracy of the position of indels (Löytynoja and Goldman 2008; Laurie et al. 2012; Villanueva-Cañas et al. 2013). The amino acid alignments were converted to coding sequence alignments using PAL2NAL (Suyama et al. 2006).

### Construction of plasmids

Full length cDNA clones for human *LHX2* and *LHX9* (IMAGE clones: IRATp970G12116D and IRCMp5012C0737Q, respectively) and *PHOX2A* and *PHOX2B* (IRAUp969B08103D and IRAUp969C0347D, respectively) were purchased from ImaGenes and checked by sequencing. The open reading frames were amplified by polymerase chain reaction with specific primers (supplementary table S13, Supplementary Material online) and they were inserted in-frame into the EcoRI and XbaI sites of the expression plasmid pGal4-DBD (de la Luna et al. 1999) to generate N-terminally fused proteins to Gal4 DBD. The deletion of the LHX2 LCR -AAAAAAAAAAKSAGLGAAGA-was performed by site-directed mutagenesis (Stratagene) on pG4DBD-LHX2 with primer LHX2ΔLCR to generate pG4DBD-LHX2ΔLCR. A synthetic gene containing the human PHOX2B open reading frame without the LCR - GGAAAAAAAAAAAAAAAAAAAAGGLAAAGG- was purchased from GeneArt (Life Technologies) and cloned into pGal4-DBD. All the plasmids were checked by DNA sequencing.

### Cell culture and transfection

HeLa cells were obtained from the American Type Cell Culture Collection. Cells were maintained at 37°C in DMEM supplemented with 10% fetal calf serum (FCS) and antibiotics. Transient transfections were performed using the calcium phosphate method and cells were processed 48 h after transfection.

### Immunofluorescence

Cells grown in coverslips were fixed in 4% paraformaldehyde in phosphate-buffered saline (PBS) for 15 min, permeabilized in 0.1% Triton X-100 in PBS for 10 min and blocked with 10% FCS for at least 30 min. Cells were incubated with a mouse monoclonal antibody to Gal4-DBD (1:100 in PBS-1% FCS; Santa Cruz Biotechnology) as the primary antibody for 1 h. After washing extensively with PBS-1% FCS, the coverslips were incubated with an Alexa Fluor 488 anti-mouse antibody (1:2000 in PBS-1% FCS; Invitrogen) for 1 h, washed repeatedly with PBS-1% FCS and mounted onto slides using Mowiol-DAPI (4′, 6′-diamino-2-phenylindole, 1 g/ml; Vector Laboratories, Inc). All procedures were done at room temperature. Images were captured with an Inverted ZEISS Microscope with fluorescence.

### Western blot

Whole-cell extracts were prepared in 25 mM Tris-HCl, pH 7.5, 1 mM EDTA, and 1% sodium dodecyl sulfate (SDS). Samples were resolved by SDS-PAGE, transferred onto nitrocellulose membrane (Hybond C; GE Healthcare), and blocked with 10% skimmed milk in Tris-buffered saline (TBS) (10 mM Tris-HCl, pH 7.5, and 100 mM NaCl) containing 0.1% Tween 20 (TBS-T). Membranes were incubated with anti-Gal4-DBD antibody (in 5% skimmed milk in TBS-T) overnight at 4°C. After washing with TBS-T, membranes were incubated for 1 h at room temperature with horseradish peroxidase-conjugated polyclonal rabbit anti-mouse antibody (in 5% skimmed milk in TBS-T) and then washed again with TBS-T. Proteins were detected by chemiluminiscence with Western Lightning Plus-ECL (Perkin Elmer) in a LAS-3000 image analyzer (Fuji PhotoFilm).

### One-hybrid gene reporter assays

HeLa cells were transfected with the pG5E1B-luc reporter (de la Luna et al. 1999) in which *Firefly* luciferase expression is driven by five repeats of yeast Gal4-binding sites introduced upstream of the minimal adenovirus E1B promoter, together with the plasmids to express unfused Gal4-DBD, or different Gal4-DBD fusion proteins as indicated in the Figure Legends. A *Renilla* luciferase plasmid (pCMV-RNL, Promega) was used as an internal control for transfection efficiency. Cells were lysed 48 h post-transfection and the activity of both luciferase enzymes was measured with the Dual-Luciferase Reporter Assay kit (Promega). Transfections were done in triplicates, and the experiments repeated independently three times.

## Supplementary Material Online

Supplementary tables are in Rado-Trilla_etal_2015_sup.xlsx, available from evolutionarygenomics.imim.es, Publications/Datasets links.

## Acknowledgements

We received funding from Ministerio de Innovación y Tecnología (BIO2009-08160 to M.M.A.), Ministerio de Economía y Competitividad (BFU2012-36820 to M.A. and BFU2010-15347 and ‘Centro de Excelencia Severo Ochoa 2013-2017′-SEV-2012-0208 to S.L.), the Secretariat of Universities and Research-Government of Catalonia (2014SGR1121 TO M.M.A. and 2014SGR674 to S.L.), Fundació Javier Lamas Miralles (fellowship to N.R-T.), and Institució Catalana de Recerca i Estudis Avançats (ICREA contract to M.M.A. and S.L.). We acknowledge Will Blevins for his useful comments on the manuscript.

